# Chemical Proteomic Profiling of the Histaminylation Proteome in Cancer Cells Unveils Uncharted Epigenetic Marks on Core Histones

**DOI:** 10.64898/2026.03.07.710331

**Authors:** Xingyu Ma, Anne A. Leaman, Zeng Lin, Huapeng Li, Zhengjun Cai, Qianyue Wang, Kaiwaan Dalal, Md Shahadat Hossain, Venkatesh P. Thirumalaikumar, Zhihong Wang, Valerie P. O’Brien, W. Andy Tao, Qingfei Zheng

## Abstract

Histamine is a key signaling molecule in pathophysiology that can exhibit significant regulatory roles in diverse health and disease status. Besides the well-studied noncovalent interactions between histamine and its receptors, protein histaminylation is a recently discovered mechanism of action, through which histamine regulates cellular signaling pathways in a covalent modification manner. Histaminylation is an emerging protein post-translational modification, where an isopeptide bond is formed between the histamine primary amine and γ-carboxyl group of glutamine through a transamidation reaction catalyzed by transglutaminase 2 (TGM2). However, due to the lack of efficient pan-specific antibodies targeting histaminylated glutamine, the histaminylation proteome in cells remains poorly explored. Here, we report the design and development of a novel N^τ^-propargylated histamine (N^τ^-PH) probe as well as its successful application in chemical proteomic profiling of the histaminylation proteome in cancer cells. Notably, new TGM2-catalyzed epigenetic marks on core histones, *e*.*g*., H2AX-Q84 and Q104 histaminylation, have been identified from cancer cells and verified. Lastly, the crosstalk between H2AX histaminylation and γH2AX formation was discovered in this study, suggesting that TGM2-mediated histaminylation plays a critical role in DNA damage responses.

## Introduction

Discovered from vertebrate tissues almost a century ago,^1^ histamine has been known as an essential hormone and neurotransmitter molecule that exhibits significant regulatory functions in pathophysiological states, such as immune defense, allergic response, digestion, brain functions, cancer development, *etc*.^2,3^ Previous studies have shown that histamine exerts its biological effects primarily by binding to G protein-coupled histamine receptors, *i*.*e*., H1, H2, H3, and H4,^4^ as well as activating ligand-gated chloride channels in the brain and intestinal epithelium.^5^ Through these noncovalent interactions with the corresponding membrane proteins, histamine became one of the earliest identified mediators of allergy, autoimmune conditions, gastric acid secretion, and hematopoiesis.^6^ Due to histamine’s high clinical relevance, a variety of drugs have been developed to target histamine-involved signaling pathways and are widely used in clinical practice as antihistamines.^7^ For example, second-generation antihistamines (*e*.*g*., loratadine, cetirizine, and fexofenadine) are commonly applied in the clinic to relieve symptoms associated with allergies and hives.^8^

In addition to noncovalent interactions with receptor proteins, recent studies have illustrated a novel covalent bioconjugation between histamine and the target protein through the formation of an isopeptide bond.^9-11^ This unique protein post-translational modification (PTM), termed histaminylation, is catalyzed by human transglutaminase 2 (TGM2) via a reversible transamidation reaction, where an amide bond is formed between the histamine primary amine and γ-carboxyl group of glutamine side chain (**Figure 1**). In our previous studies, we for the first time identified histaminylation occurring on histone H3 glutamine 5 (H3-Q5), a new epigenetic mark that plays a role in circadian rhythm regulation.^9^ Importantly, we discovered that the dynamics (including installation, removal, and replacement) of H3-Q5 histaminylation (H3-Q5his) are regulated by a single enzyme, *i*.*e*., TGM2.^9^ Unlike H3-Q5 serotonylation (H3-Q5ser) that can form a cation-π interaction with WD repeat domain 5 (WDR5) to enhance its binding to H3, positively charged H3-Q5his inhibits the binding affinity between the H3 N-terminal tail and the WDR5 subunit of H3K4 methyltransferase complexes (MLL1-4 and SETD1A/B) through charge repulsion, thereby antagonizing H3K4 methylation.^9-12^ These findings provide a new mechanism for histamine’s significant biological effects in cell signaling, where histamine interacts with its target proteins in a covalent manner via modifying the corresponding glutamine residues.

**Figure 1.**
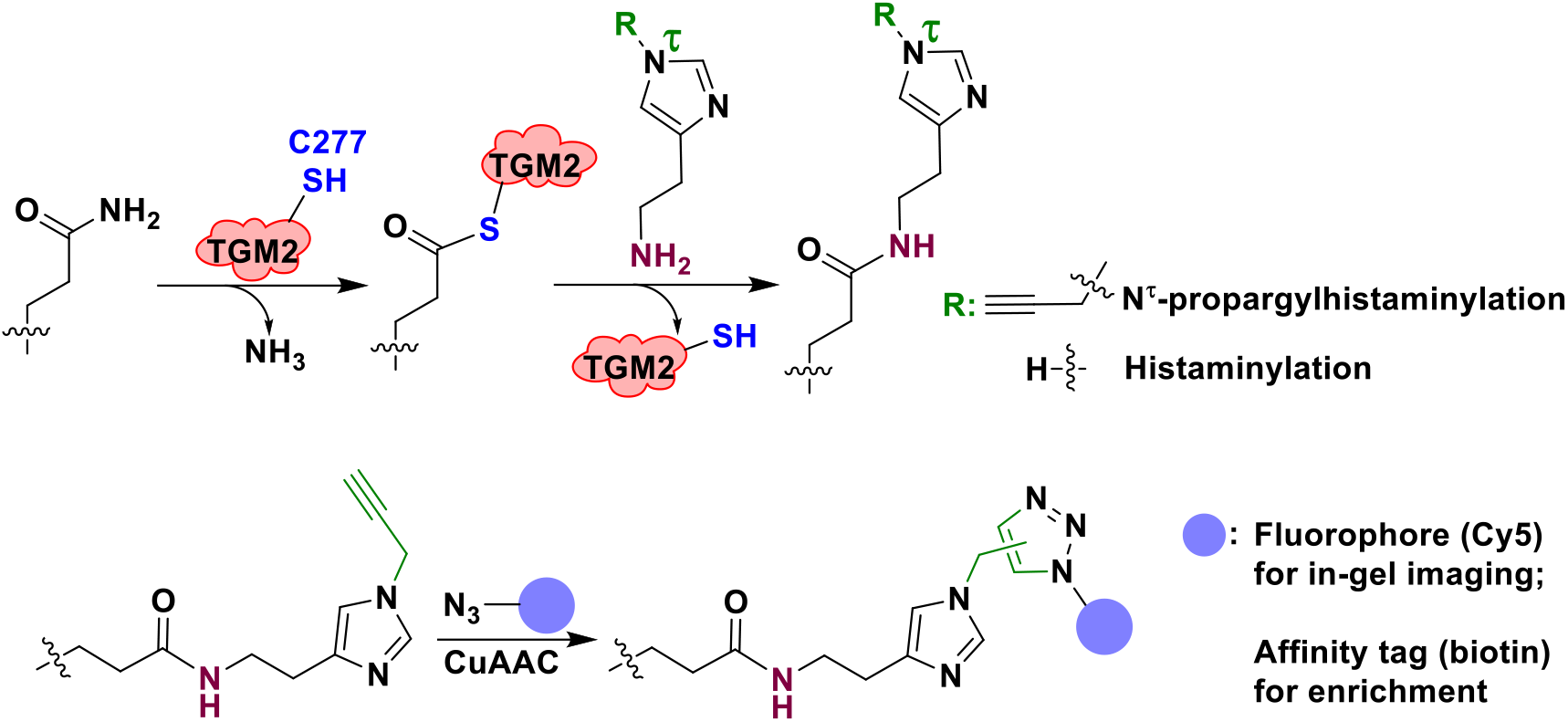
The transglutaminase 2 (TGM2)-mediated histaminylation reaction, which is dependent on its catalytic cysteine (C277), and the structure of the N^τ^-propargylated histamine (N^τ^-PH) probe used in this study for in-gel imaging and chemical proteomics.

Histone H3 was not the first histaminylation target identified, and in fact other proteins (especially G proteins) that undergo TGM2-mediated histaminylation have been known for decades.^13^ With that being said, the global profiling of the cellular histaminylation proteome has not been successfully performed, due to the lack of efficient molecular tools (such as pan-specific antibodies targeting histaminylated glutamine). Here, we designed and synthesized a novel N^τ^-propargylated histamine (N^τ^-PH) probe with a click handle that can be well recognized by TGM2 and efficiently incorporated onto the glutamine residues of target proteins for further labeling and enrichment (**Figure 1**). Utilizing this powerful chemical biology probe, we conducted a chemoproteomic analysis of histaminylation proteins in colorectal and stomach cancer cell lines that have a high expression level of endogenous TGM2.^14,15^ Therefore, many histaminylated proteins and their corresponding PTM site were identified. Unexpectedly, several uncharted epigenetic marks on core histones (*e*.*g*., H2AX-Q84his and Q104his) have been discovered from cancer cells and verified via biochemical assays.

## Results and Discussion

### Design and Synthesis of the N^**τ**^-PH Probe for *in vitro* Studies of TGM2-mediated Histaminylation

First, to specifically label and enrich histaminylated proteins, a click handle-containing probe, N^τ^-PH, was synthesized in three steps (**Figure 2A**), of which the propargyl group on the N^τ^ atom of the imidazole ring cannot perturb the transamidation reaction catalyzed by TGM2. Correspondingly, N^α^-PH was synthesized as a negative control probe, which possesses the same molecular weight as N^τ^-PH but cannot undergo transamidation mediated by TGM2 (**Figures 2A and 2B**). The structures of N^τ^-PH and N^α^-PH together with their synthetic intermediates were confirmed using ^1^H and ^13^C nuclear magnetic resonance (NMR) analyses (**Figure S1**). A twelve-amino-acid (12-aa) peptide of the histone H3 N-terminal tail that contains the known histaminylation site (H3-Q5) was utilized as a substrate to test the reactivity of N^τ^-PH in the TGM2-catalyzed transamidation reaction. Matrix-assisted laser desorption/ionization-time-of-flight mass spectrometry (MALDI-TOF MS) analysis showed that N^τ^-PH can be well recognized by TGM2 and its reactivity as a monoaminylation donor in the transamidation reaction is comparable to histamine (**Figure 2C**).

**Figure 2.**
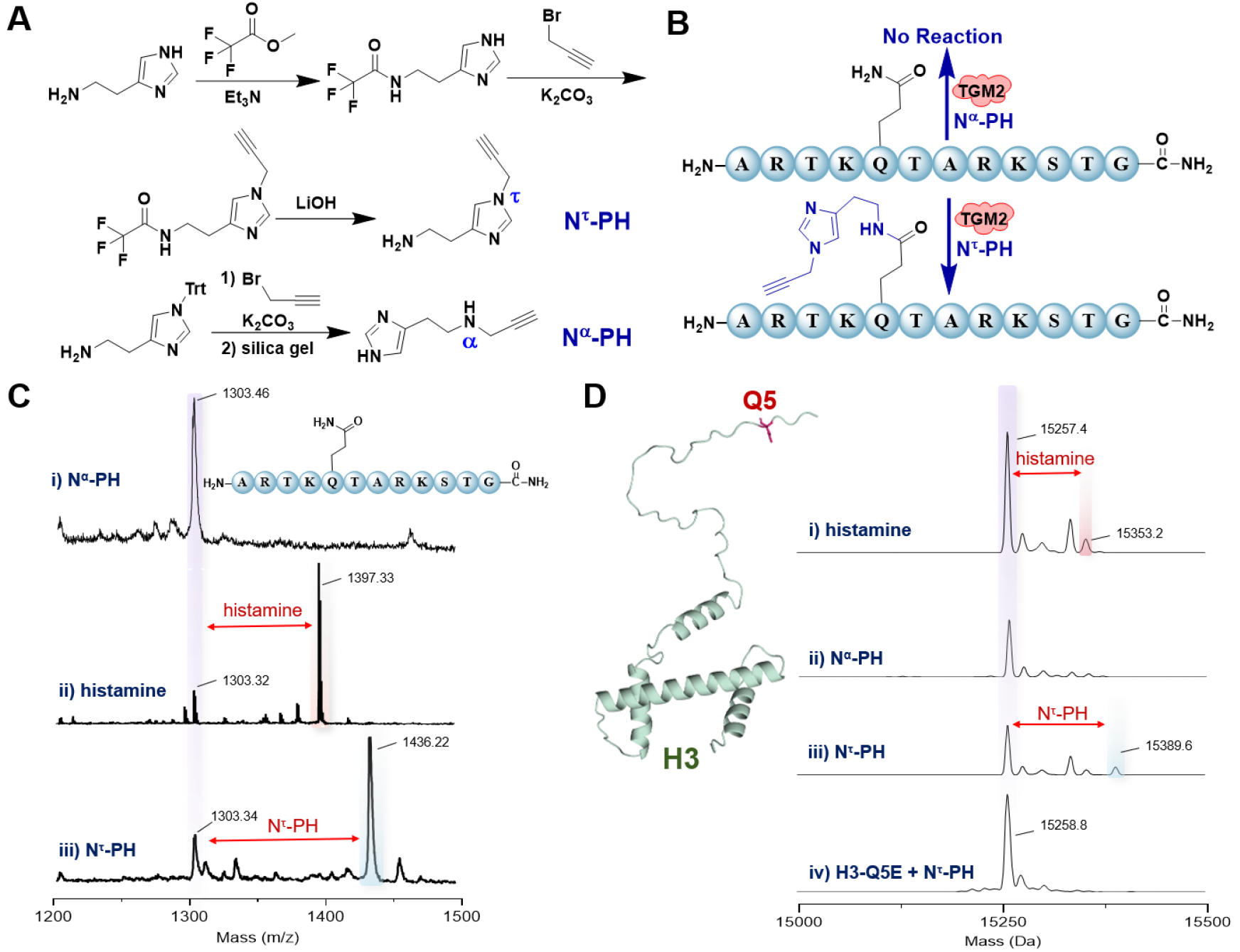
Synthesis of N^τ^-PH as a chemical probe to study histaminylation and N^α^-PH as a negative control. (**A**) Synthetic routes of N^τ^-PH and N^α^-PH. (**B**) Reactivity and (**C**) MALDI-TOF MS analysis of N^τ^-PH (or N^α^-PH) against H3 N-terminal tail peptide (1-12 aa). The peptide substrate was incubated with TGM2 and (i) N^α^-PH, (ii) histamine, or (iii) N^τ^-PH, respectively. (**D**) LC-MS (Sciex ZenoTOF 7600) analysis of TGM2-mediated transamidation between histamine derivatives and full-length histone H3 variants after deconvolution. The *in vitro* biochemical reactions are between (**i**) wild-type H3 and histamine, (**ii**) wild-type H3 and N^α^-PH, (**iii**) wild-type H3 and N^τ^-PH, and (**iv**) H3-Q5E mutant and N^τ^-PH.

Thereafter, full-length H3 proteins (wild-type H3 and H3-Q5E mutant, which cannot be histaminylated and serves as a negative control) were used as the substrates, and liquid chromatography-mass spectrometry (LC-MS) analysis suggested that H3-Q5 is the PTM site induced by N^τ^-PH, which is in line with the reactions with histamine, while N^α^-PH could not be installed onto the corresponding target site (**Figure 2B**). The following LC-MS-based enzyme kinetics analyses further confirmed that N^τ^-PH exhibits the similar reactivity to histamine in TGM2-catalyzed H3-Q5 monoaminylation, acting as an ideal competitor probe for studying protein histaminylation (**Figure S2**). Overall, the substitution of a N^τ^-H bond within histamine’s imidazole ring by a propargyl group does not influence its nucleophilic property as a monoaminylation donor in TGM2-catalyzed protein PTM. Thus, the alkynyl group-possessing probe, N^τ^-PH, can serve as a mimic of histamine for the labeling and enrichment of histaminylated proteins via copper(I)-catalyzed azide-alkyne cycloaddition (CuAAC).^16^

### Applications of the N^**τ**^-PH Probe for Chemical Proteomics and Fluorescence Imaging Analysis of Histaminylation in Cancer Cells

Utilized as an alternative to a pan-specific antibody, the N^τ^-PH probe was incubated with a colorectal cancer cell line, HCT 116, and incorporated into the target proteins by endogenously expressed TGM2. Azid-Cy5 was used as an indicator for the in-gel imaging assays, which can be conjugated onto N^τ^-PH-modified glutamine residues of the histaminylation proteome through CuAAC (**Figures 1** and **3A**). The in-gel imaging assays showed that N^τ^-PH is cell-permeable and stable in mammalian cells. The dose-dependent assay and competition treatment of histamine as well as other monoamines (*i*.*e*., serotonin and dopamine)^17,18^ further proved that N^τ^-PH is an ideal mimic of histamine as a donor for protein monoaminylation (**Figure S3**). These findings are consistent with the enzyme kinetics analyses, which revealed that TGM2 exhibits a markedly higher preference for histamine and N^τ^-PH than for dopamine or serotonin as transamidation substrates (**Figure S2**). Given the utility of the N^τ^-PH probe for incorporation into the modified proteins by TGM2 as a histamine mimic, we treated HCT 116 cells with N^τ^-PH and performed CuAAC-based chemical proteomics analysis using an azide-tagged biotin for enrichment. Volcano plots showed that over 450 proteins were enriched in the probe-treated group and modified by N^τ^-PH on their corresponding glutamine residues (**Figure 3A**). Similarly, a stomach cancer cell line, AGS, was used for N^τ^-PH-based chemoproteomic analysis, where over 200 proteins were identified to possess histaminylation. 104 proteins were modified in both HCT 116 and AGS cells (**Figure S4**).

**Figure 3.**
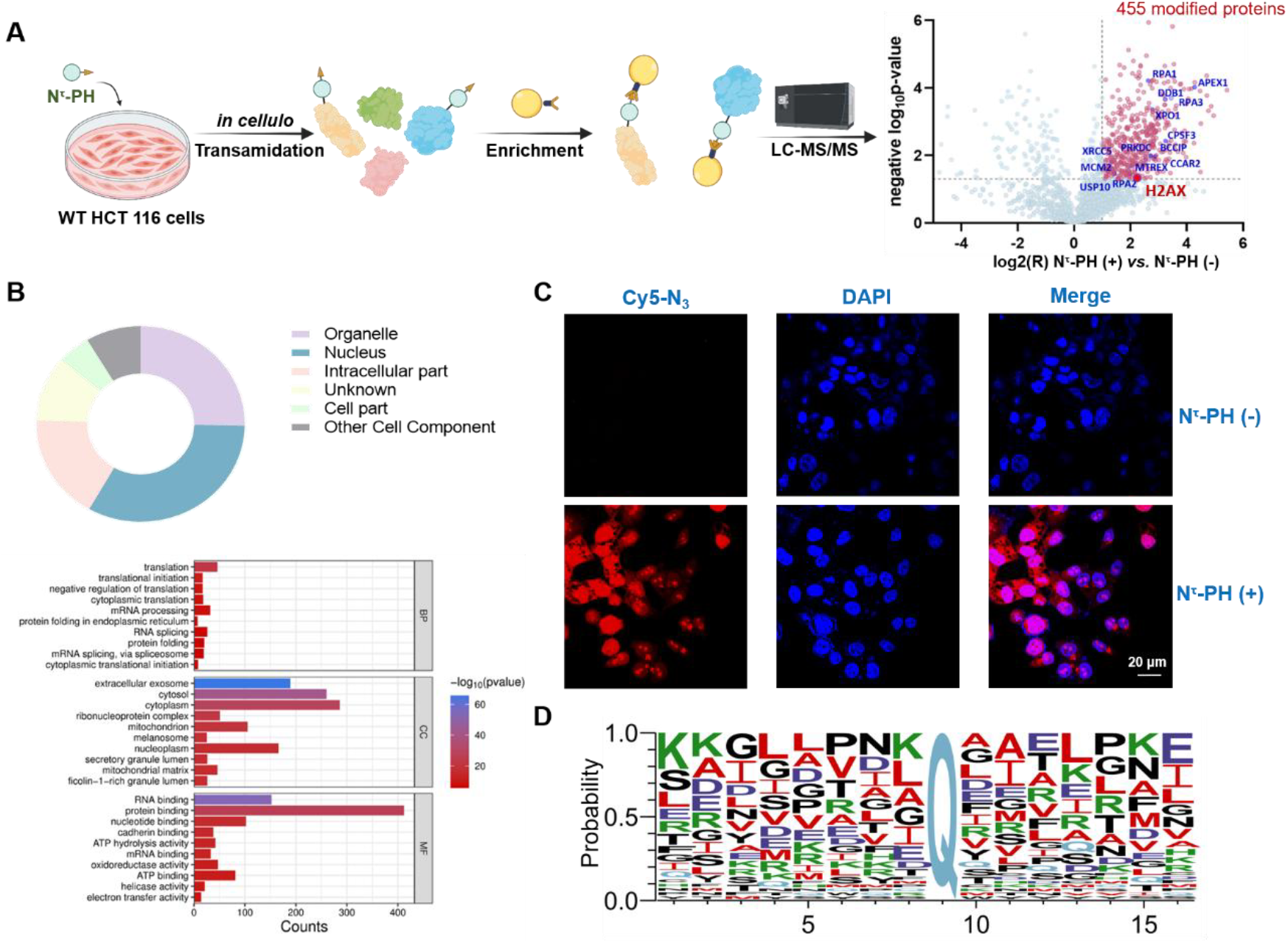
Chemical proteomic profiling of the histaminylation proteome in wild-type (WT) HCT 116 cells. (**A**) Workflow of N^τ^-PH probe-based chemical proteomics analysis of histaminylated proteins in HCT 116 cells, where 455 modified proteins were identified. The modified proteins that are involved in DDR pathways were annotated. (**B**) Bioinformatic analysis of the identified proteins in this study showing the subcellular locations and functions of the histaminylation proteome in cancer cells. (**C**) Confocal microscopy analysis using the N^τ^-PH probe shows histaminylated proteins are mainly nucleoproteins. (**D**) pLogo analysis of the identified modification sites in this study indicates TGM2 has a broad substrate specificity.

In line with our previous discoveries,^19,20^ bioinformatics analysis showed that nucleoproteins, which play critical roles in gene transcription regulation and cancer development, occupied a large portion of the histaminylated proteome (**Figure 3B**). Confocal microscopy analysis using the N^τ^-PH probe and CuAAC for imaging further confirmed the subcellular locations of histaminylated proteins within cancer cells (**Figure 3C**). In addition, many histaminylated proteins identified in the N^τ^-PH probe-based proteomics are involved in DNA damage response (DDR) pathways (*e*.*g*., H2AX, DDB1, RPA1, BCCIP, APEX1, *etc*),^21^ suggesting a potential crosstalk between protein histaminylation and DDR in caner development.

A cleavable linker (*i*.*e*., dialkoxydiphenylsilane)-containing biotin azide was thereafter employed in the chemical proteomics experiments for PTM site analysis. Numerous histaminylation sites were thus identified (**Figure S5**). Several histaminylation sites were verified by using the *in vitro* enzymatic assays (**Figure S6**), including HSP90AB1-Q80, ACTA2-Q248, KRT6C-Q401, RYR3-Q1316, and H2AX-Q84/Q104. Notably, pLogo analysis of the histaminylated peptide sequences indicated that TGM2 has a high enzyme promiscuity, without an obviously preferred substrate sequence (**Figure 3D**).

### Identification of Uncharted Histaminylation Sites on Core Histones using the N^**τ**^-PH Probe

Among the identified histaminylation sites, the two glutamine residues within histone H2AX (*i*.*e*., Q84 in the globular domain and Q104 in the unstructured C-terminus) particularly piqued our interest (**Figures 4A and 4B**). Even though *in vitro* biochemical reactions revealed that the two glutamine residues Q123 and Q124 within the unstructured C-terminal tails of histones H2AZ and H2AV (not present in canonical H2A) were also the monoaminylation (*i*.*e*., serotonylation) sites catalyzed by TGM2,^22^ currently H3-Q5 is the primary histaminylation site on histones that was identified from cells/tissues and verified *in vitro*/*in vivo*, the epigenetic function of which has been well established.^9-11^ Given the two new histaminylation sites on H2AX identified using our chemical proteomics approach, we synthesized two 11-aa peptides to confirm H2AX-Q84 and Q104 are indeed substrates of TGM2-catalyzed histaminylation. *In vitro* biochemical assays illustrated that TGM2 could efficiently install histamine onto H2AX-Q84 and Q104, while the catalytically inactive mutant, TGM2-C277A, could not (**Figure S5**). Thereafter, full-length H2AX was purified recombinantly from *E. coli* and treated with TGM2 and N^τ^-PH or N^α^-PH. CuAAC-based in-gel imaging and immune blotting further confirmed that N^τ^-PH could be efficiently incorporated into H2AX by TGM2 through enzymatic transamidation (**Figure 4C**).

**Figure 4.**
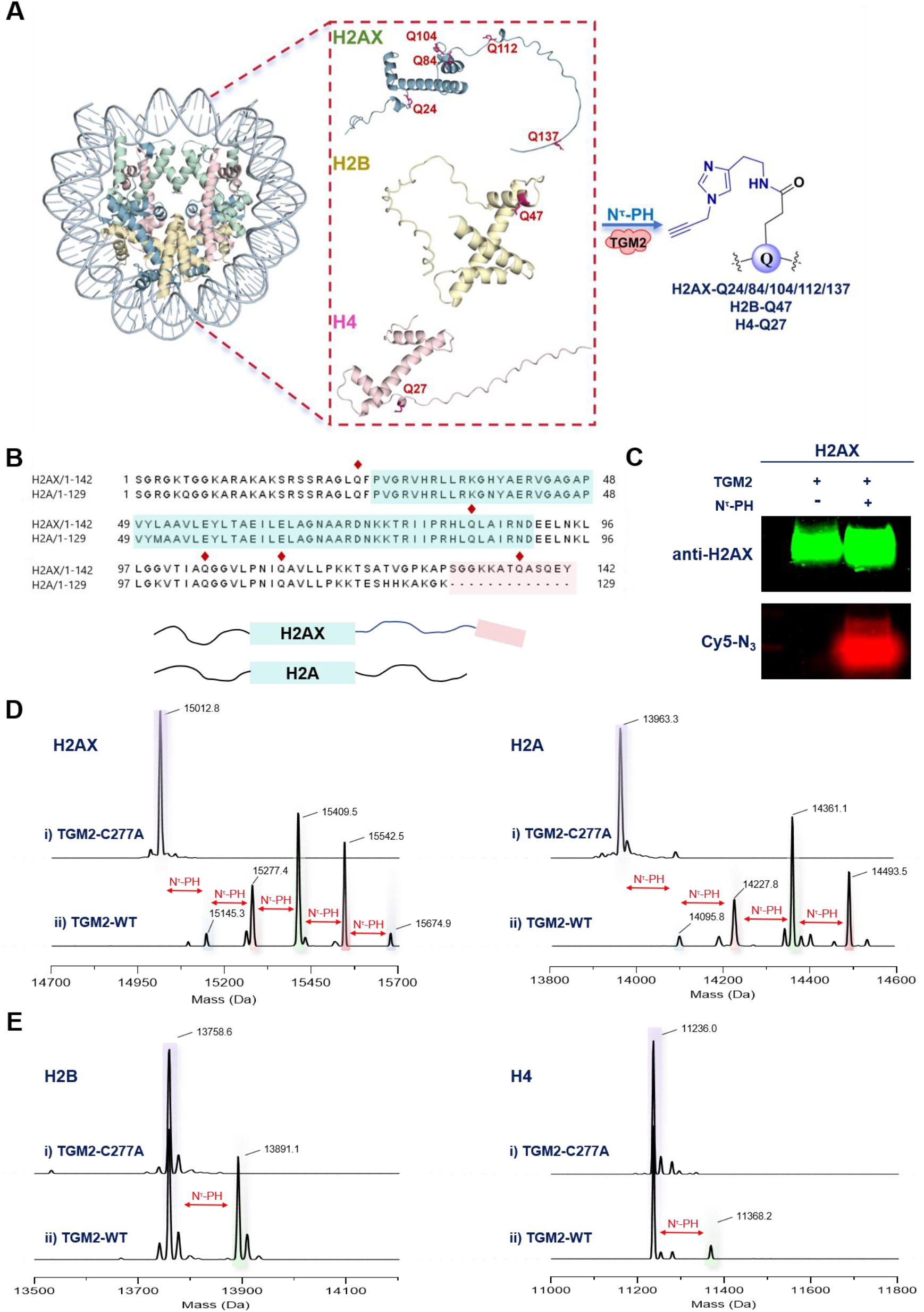
Identification of seven new histaminylation sites on core histones as uncharted epigenetic marks. (**A**) Structure of H2AX-containing nucleosome (PDB code: 9MMK) and the new histaminylation sites identified on H2AX (alphafold code: AF-P16104-F1-v6), H2B, and H4. (**B**) Sequence alignment of H2AX and H2A, where the globular domain is indicated in a light green box and the unique unstructured C-terminal tail of H2AX is highlighted in a pink box. The identified glutamine residues that undergo histaminylation are labeled by red diamonds. (**C**) CuAAC-based in-gel imaging and immunoblotting using N^τ^-PH as the chemical probe shows full-length H2AX can be histaminylated by TGM2 through transamidation. (**D**) LC-MS (Sciex ZenoTOF 7600) analysis of TGM2-catalyzed transamidation reactions after deconvolution, showing there are five modified glutamines in H2AX (Q24/84/104/112/137) and four modified glutamines in H2A (Q24/84/104/112). (**E**) LC-MS (Sciex ZenoTOF 7600) analysis of TGM2-catalyzed transamidation reactions after deconvolution, showing there is one modified glutamine in H2B (Q47) and one modified glutamine in H4 (Q27). For the *in vitro* biochemical reactions, the substrate was incubated with N^τ^-PH and (**i**) wild-type TGM2 (TGM2-WT) or (**ii**) its catalytically inactive mutant (TGM2-C277A).

Notably, the sequences of these two peptide substrates are conserved in canonical histone H2A (**Figure 4B**). Thus, recombinant nucleosome core particles (NCPs) containing full-length H2AX or H2A were compared as the substrates for *in vitro* histaminylation tests, where N^τ^-PH was utilized as the monoamine donor. Unexpectedly, LC-MS analyses showed that four histaminylation sites were introduced by TGM2 onto H2A, while H2AX was modified by five N^τ^-PH probes (**Figure 4D**). As there are four conserved glutamine residues within the H2AX and H2A sequences (**Figure 4B**), our results suggested that one more glutamine within its unique C-terminal tail was histaminylated by TGM2. Therefore, bottom-up mass spectrometry was performed to analyze the PTM sites of histaminylated H2AX and H2A and the corresponding result illustrated that the modified glutamine residues are Q24, Q84, Q104, Q112 of H2AX and H2A as well as Q137 of H2AX (**Figures S7** and **S8**). More intriguingly, H2B-Q47 and H4-Q27 were also found to be histaminylated in the NCP-based *in vitro* biochemical reaction (**Figures 4E, S9**, and **S10**). The bottom-up MS analysis of histone fractions extracted from N^τ^-PH-treated HCT 116 cells revealed H2A/H2AX-Q112 as a histaminylation site *in vivo* (**Figure S11**). Moreover, a recent study reported H4-Q27 as a TGM2-mediated monoaminylation site in cancer cells.^23^ Overall, these results provided direct *in vitro* evidence for the first time that in addition to the reported histaminylation site on H3 (*i*.*e*., H3-Q5his),^9^ TGM2-mediated H2AX-Q24his/Q84his/Q104his/Q112his/Q137his, H2B-Q47his, and H4-Q27his are seven newly discovered epigenetic marks that may exhibit epigenetic effects of importance (**Figures 4A**).

### Discovery of the Unknown Crosstalk between H2AX histaminylation and γH2AX using the N^**τ**^-PH Probe

To investigate the crosstalk between H2AX histaminylation and γH2AX formation (*i*.*e*., H2AX-S139 phosphorylation), which is a key biomarker in DNA double-strand break (DSB) repair,^24^ we constructed the CRISPR-mediated TGM2 knockout (TGM2-KO) HCT 116 cell line using the same vectors in our previous study.^9^ N^τ^-PH-based chemical proteomics and confocal imaging demonstrated that the amount of histaminylated proteins within the TGM2-KO cells was significantly downregulated compared with the wild-type (WT) HCT 116 cells (dropped from 455 to 27), including H2AX (**Figures 5A** and **5B**). DSBs in WT and TGM2-KO cells treated with N^τ^-PH were induced using ultraviolet (UV) radiation and H_2_O_2_ treatment, and the histone factions were subsequently extracted (**Figure 5C**). Antibody- and N^τ^-PH-based immune blotting illustrated TGM2 elevates the levels of both H2AX histaminylation and γH2AX, suggesting a previously unknown crosstalk between these two epigenetic marks within H2AX (**Figure 5D**). Notably, the deficiency of TGM2 and protein histaminylation in HCT 116 cells significantly decreases the cell viability (**Figure S12**), which is consistent with the positive role γH2AX plays in gene transcription and cell survival.^25^ These results also suggest that targeting TGM2-mediated H2AX histaminylation and the associated DDR pathways may serve as a novel anti-cancer strategy.

**Figure 5.**
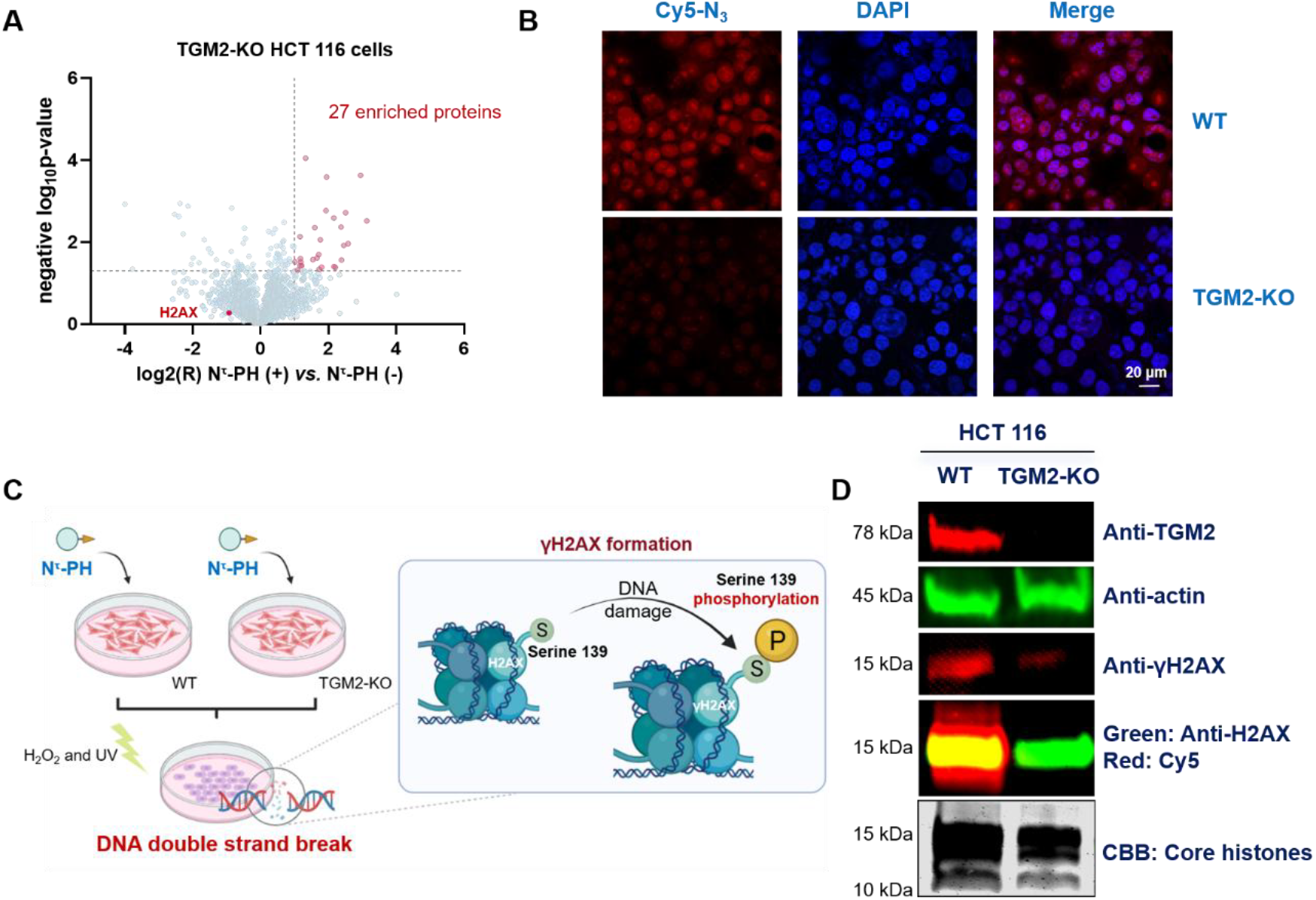
Validation of the crosstalk between TGM2-mediated H2AX histaminylation and γH2AX formation. (**A**) Chemical proteomics analysis of the histaminylated proteins in TGM2-KO HCT 116 cells using the N^τ^-PH probe. (**B**) Confocal microscopy analysis of the histaminylation proteome using the N^τ^-PH probe in WT and TGM2-KO HCT 116 cells. (**C**) Schematic workflow for inducing DNA double-strand breaks (DSBs) and γH2AX formation in N^τ^-PH-treated WT and TGM2-KO HCT 116 cells. CuAAC-based in-gel fluorescence imaging and immunoblotting analysis using N^τ^-PH probe, unveiling the crosstalk between H2AX histaminylation and γH2AX. The cytoplasmic protein and histone fractions were extracted separately from WT and TGM2-KO HCT 116 cells and analyzed using the corresponding antibodies or CuAAC-based fluorescence imaging. The Cy5 signal indicated H2AX histaminylation levels. Coomassie Brilliant Blue (CBB) staining of core histones was used as the loading control for analyzing the histone fractions.

## Conclusions

Histamine has long been known as an essential signaling molecule that is involved in diverse cellular pathways.^1-6^ As an emerging protein PTM, TGM2-mediated histaminylation offers a new mechanism for histamine’s regulatory functions in pathophysiology. In this research, we first designed and synthesized a structural analog of histamine, *i*.*e*., N^τ^-PH, and proved it was an ideal substrate mimicking histamine in TGM2-catalyzed transamidation reactions. Thereafter, chemical proteomic analysis was conducted using this chemical biology probe and over five hundred proteins containing histaminylation sites were identified in HCT 116 and AGS cancer cells. Several modification sites (including HSP90AB1-Q80, ACTA2-Q248, KRT6C-Q401, RYR3-Q1316, *etc*.) were validated via *in vitro* enzymatic assays.

Before this work, H3-Q5 was the only histaminylation site identified in cells and was shown to exhibit significant epigenetic effects in transcription regulation. Here, applying our novel chemical biology tools, we successfully identified and verified five new histaminylation sites in H2A variants, one new histaminylation site in H2B, and one histaminylation site in H4 as uncharted epigenetic marks (**Figure 4**). Although an unknown crosstalk between TGM2-mediated H2AX histaminylation and γH2AX formation was reported in this research, further studies are needed to understand the roles of these new epigenetic marks in disease development, especially cancer. Overall, utilizing cutting-edge chemical biology approaches, in this study we for the first time reported the histaminylation proteome in cancer cells and unveiled several unreported epigenetic marks on core histones, which provide new insights to understand the diversity and functionality of protein histaminylation and open a door toward future cancer therapeutics by targeting TGM2-mediated monoaminylations.^14,15^

## Supporting Information

The Supporting Information is available free of charge at https://pubs.acs.org/doi/xxxxxxx.

## Acknowledgements

This research was financially supported by the NIH (R35 GM150676) and startup funds from Purdue University (for Q.Z.). A.W.T. was supported by NSF funding (CHE-2404098).

## For Table of Contents use only

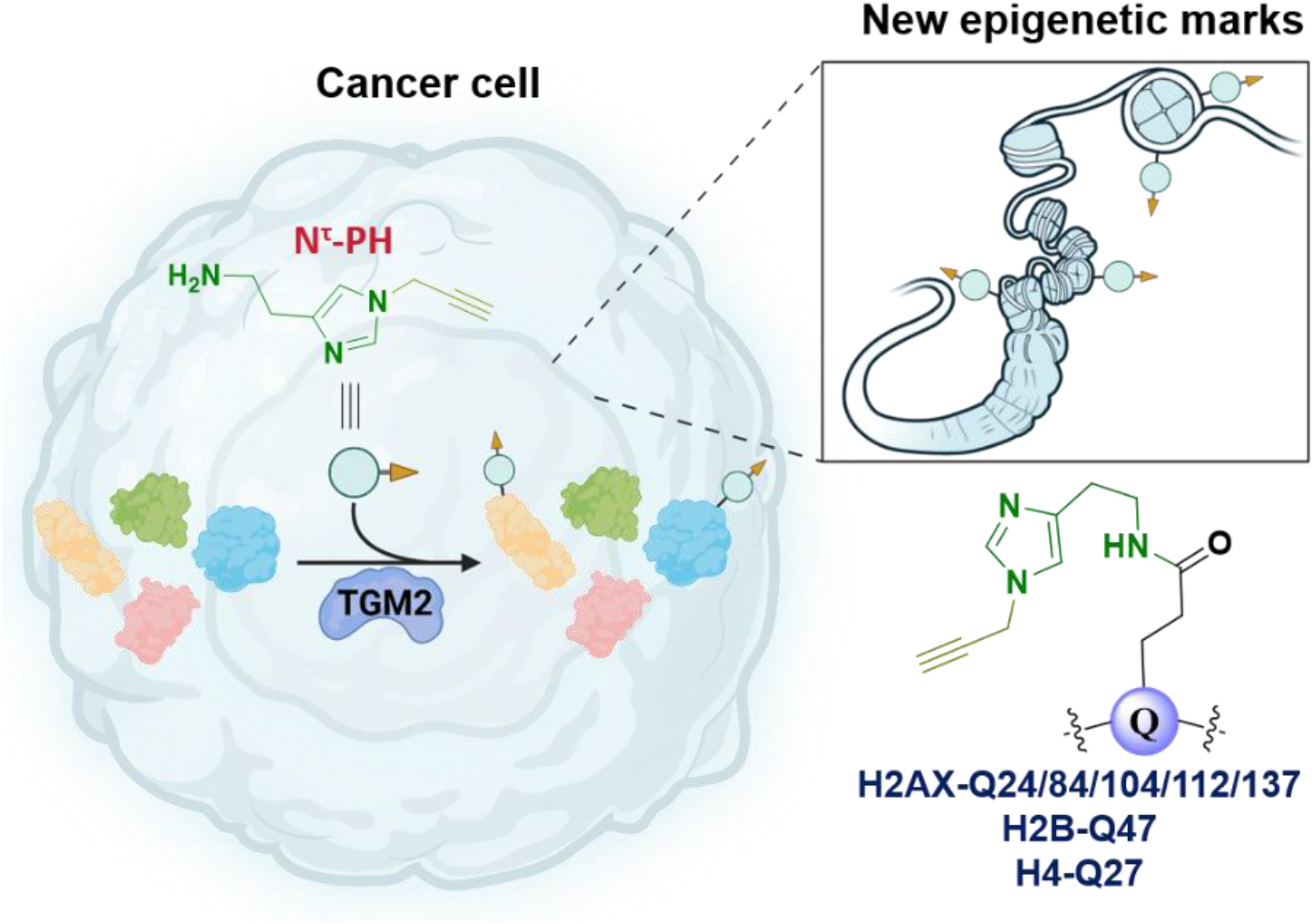

